# Selective influence of dopamine on electrocortical signatures of error monitoring: a combined EEG and immersive virtual reality study in Parkinson’s disease

**DOI:** 10.1101/2022.02.05.478638

**Authors:** R. Pezzetta, D.G. Ozkan, V. Era, G. Tieri, S. Zabberoni, S. Taglieri, A. Costa, A. Peppe, C. Caltagirone, S.M. Aglioti

## Abstract

Detecting errors in one’s own and other’s actions is likely linked to the discrepancy between intended or expected and produced or observed output. To detect and process the occurrence of salient events seems associated to the release of dopamine, the balance of which is profoundly altered in Parkinson’s disease (PD). EEG studies in healthy participants indicate that the occurrence of errors in observed actions triggers a variety of electrocortical indices (like mid-frontal theta activity, error-related delta and the Error Positivity, oPe), that seem to map different aspects of error detection and performance monitoring. Whether these indices are differently modulated by dopamine in the same individual has never been investigated. To explore this issue, we recorded EEG markers of error detection by asking healthy controls (HCs) and PD patients to observe ecological reach-to-grasp a glass actions performed by a virtual arm seen in first person perspective. PD patients were tested under their dopaminergic medication (‘on-condition’), and after dopaminergic withdrawal (‘off-condition’). HCs showed a clear oPe and an increase of delta and theta power during the observation of erroneous vs. correct actions. In PD patients, oPe and delta responses were always preserved. Crucially, however, an error-related increase of theta power was found in ‘on’ but not in ‘off’ state PD patients. Thus, different EEG error signatures may index the activity of independent systems and error related theta power is selectively modulated by dopamine depletion. Our findings may pave the way to the discovery of dopamine-related biomarkers of higher-order motor cognition dysfunctions that may have crucial theoretical and clinical implications.

**Significance Statement:** Dopaminergic neurons respond to salient events during performance monitoring. Yet, the impact of dopamine depletion on the human reactivity to observed errors is still unclear. We recorded EEG in patients with Parkinson’s Disease (PD) under dopaminergic treatment (‘on-condition’) and medication withdrawal (‘off-condition’) while they observed correct and erroneous goal-related actions performed by a virtual limb. Analysis of Error Positivity (oPe) and theta and delta power increase, markers of physiological error-monitoring, indicates that while the formers were intact, the latter was preserved in the ‘on’ and altered in the ‘off’ condition. Thus, different EEG markers of error monitoring likely rely on independent circuits. Moreover, mid-frontal theta activity alterations may represent a marker of dopamine-related neurophysiological impairments of higher-order cognition.

## Introduction

The progressive degeneration of dopaminergic neurons in the substantia nigra pars compacta that characterizes Parkinson’s Disease (PD) brings about alterations in a complex circuit involving subcortical and cortical (mainly frontal and cingulate) regions (Parkinson, 1817; Ullsperger & Von Cramon, 2006; Wylie et al., 2010; Zavala et al., 2018) that lead not only to motor symptoms, but also to deficits of higher order cognitive functions (Chaudhuri et al., 2010; Ponsi et al., 2021), including performance monitoring (Seer et al., 2016). Studies about the influence of dopamine on cognitive functions hint at its role in regulating predictive processes (Clark, 2013; Friston & Kiebel, 2009). Indeed, dopamine is released in response to salient and unexpected events, such as unpredicted errors (Gardner, et al., 2018; Holroyd & Coles, 2002; Schultz, 1998, 2016).

Making an error triggers specific EEG signatures (Mid-frontal theta increase, increase in delta power, Error-Related negativity, Positivity Error; Cavanagh et al., 2012; Joch et al., 2017; Ridderinkhof et al., 2004a). Although, smaller in amplitude and higher in latency (Koban & Pourtois, 2014) similar signatures are also triggered by mere action error observation (de Bruijn et al., 2007). Specifically, observation of others’ errors is accompanied by increased mid-frontal theta, frontal error-related negativity (oERN – observed error related negativity), and a Positivity Error (oPe – observed positivity error) that may be seen not only when one observes an error committed by another person (van Schie et al., 2004), or by a partner during motor interactions (Era et al. 2019; Moreau et al., 2020) but also when committed by an embodied virtual arm seen in first-person perspective (1PP; Pavone et al., 2016; Spinelli et al. 2018). From a neuroanatomical and functional point of view, mounting evidence suggest that the prefrontal cortex, including the dorsal ACC generates error-related midfrontal theta/ERN as a low-level mismatch between the actual and expected response (Cavanagh et al., 2014; Jocham & Ullsperger, 2009; Parker et al., 2005; Ridderinkhof et al., 2004b). On the other hand, less clear is the origin of the Pe, which has been associated with the engagement of different areas such as the anterior parts of the ACC (Holroyd & Coles, 2002), posterior portions of the cingulate (Vocat et al., 2008) and insular cortices (Klein et al., 2013), which in turn have been associated to the salience network (Lenzoni et al., 2021; Orr et al., 2011; Wessel, 2017); the Pe/oPe seem to signal the awareness of an error, following motivational and affective events, and may be link with post-error adaptation (Overbeek et al., 2005; Danielmeier & Ullsperger, 2011). The distinct cortical involvement of these error-related responses is in line with evidence suggesting the existence of independent mechanisms underlying the error monitoring system (di Gregorio et al., 2018; Steinhauser & Yeung, 2010), that seem also to rely on distinct neurotransmitters (Falkenstein et al., 2005; Holroyd & Coles, 2002). As pointed out by Overbeek, Nieuwenhuis & Ridderinkhof (2005) and few pharmacological studies (De Bruijn et al., 2004), the midfrontal cortex is densely targeted by ascending dopaminergic projections which might affect midfrontal theta and ERN amplitude, but not the Pe and error-related delta activity, which have been associated with locus coeruleus-norepinephrine system activity (Ridderinkhof et al., 2009; Ullsperger et al., 2010; Wessel et al., 2011). On this road, studies on rats investigated the effects of neurochemical lesions of the ventral tegmental area, a major source of dopaminergic projections to the forebrain, finding a 30-46% reduction in dopamine, that however did not alter the P3-like potentials recorded intracranially (Ehlers & Chaplin, 1992; Nieuwenhuis et al., 2005). Despite a few studies pointing in this direction, the evidence is still not clear since no study has simultaneously investigated the dopaminergic influence on different error monitoring processes. In this vein, testing PD patients while under their dopaminergic medication (‘on-condition’) and after dopaminergic withdrawal (‘off-condition’) can be an ideal model for exploring the selective influence of dopamine on different EEG correlates of error monitoring. Few EEG studies reported diminished amplitude of the ERN/theta during action execution in PD, while some other studies found mixed results for the Pe (see Lenzoni et al., 2021 and Pezzetta et al., 2021 for systematic reviews). It is worth noting that only two studies used the time-frequency approach (Beste et al., 2017; Singh et al., 2018) and only one found no difference due to dopaminergic medication in conflict-related theta (Singh et al., 2018). Thus, it is still unclear whether dopamine balance is necessary for human mid-frontal theta activity during error monitoring. Beside theta, other frequencies are implied into error-monitoring processes and Parkinson’s Disease. Delta (2-4 Hz) frequencies were found associated with the Pe response, such that erroneous actions are characterized not only by the positive deflection in the time-domain but also by an enhanced delta power (Luu et al., 2004; Ullsperger et al., 2004). On the other hand, several studies found patterns of beta alteration in patients with dopaminergic deficiency (Moran et al., 2011), such as exagerated burst of beta oscillations, which has been associated with the motor impariments often observed in this population.

To investigate how dopamine balance influence error monitoring mechanisms, we recorded EEG in the same PD patients while in ‘on’ and ‘off’ condition, as well as in healthy controls. Participants were immersed in a virtual scenario and passively observed from a 1PP a virtual arm that executed correct or incorrect actions. This approach proved adept to induce the illusion of ownership over the virtual body, allowing to investigate error processing in highly realistic circumstances (Pavone et al., 2016; Pezzetta et al., 2018; Spinelli et al., 2018). Moreover, exploring action processing in the absence of overt movements allowed us to control for any confounds due the interindividual differences in task difficulty or response speed that might occur between patients.

We hypothesized that distinct and independent error processes co-exist (Di Gregorio et al., 2018), and that patients in ‘off’ condition would exhibit specific alteration of the electro-cortical markers of error processing purportedly modulated by dopamine (i.e. midfrontal theta) without affecting markers that appear to be less related to this neurotransmitter (i.e. oPe/Delta; Falkenstein et al., 2001; Luu et al., 2004). Based on the evidence that fronto-central theta is related to executive functions and working memory (Eckart et al., 2014), we also explored the relation between theta activity and tests assessing executive functions in PD. On the other hand, we expect a modulation of beta frequencies in PD, such as an increased beta activity when patients are tested without dopaminergic intake (Moran et al., 2011).

## Methods and Materials

### Participants

Seventeen patients with Parkinson Disease (PD) took part in the study. The MorePower (version 6.0.4, Campbell & Thompson, 2012) software used for computing the sample size indicated that 14 participants would be required in a design with a power of 0.85, alpha of 0.05 and a partial eta^2^ of 0.4 (as found in a previous study using the same paradigm to assess the electroencephalographic markers of error monitoring; Pezzetta et al., 2018). All the participants had normal or corrected-to-normal visual acuity. The inclusion criteria were: i) diagnosis of idiopathic PD (United Kingdom Parkinson’s Disease Society brain bank criteria, UPDRS; Huges et al., 1992); ii) absence of mental deterioration (Mini Mental State Examination, MMSE > 26); iii) absence of other neurological and psychiatric diseases; iv) treatment with daily doses of dopamine or dopamine agonists (L-Dopa equivalent doses). One patient was excluded due to probable misassumption of medication and lack of motor scale data; one patient dropped the study. Thus, a final group of 15 PD was included (5 females, 10 males; mean ± SD: Age: 70 ± 9; Years of Education: 12 ± 4). Sixteen healthy participants served as controls (HCs). One participant was excluded due to impaired vision and one to mental deterioration, thus, a group of 14 HCs - matched for age and education - was included in the study (5 females, 9 males. Mean ± SD: Age: 70 ± 6; Years of Education: 13 ± 3). HCs were included according to the following criteria: i) absence of neurological and/or psychiatric diseases in anamnesis; ii) absence of subjective cognitive disorders; iii) absence of medications with psychotropic action iv) MMSE > 26 (details in Table 1a).

**Table 1.** *Demographic and clinical data*. **a**. Summary of demographics and clinical scores for PD group and control group (HC). Age: age in years; Education: education in years; MMSE: Mini Mental State Examination; UPDRS-III: Unified Parkinson’s Disease Rating Scale section III; H &Y: Hoehn and Yahr scale. **b**. Summary of motor scale scores of PD patients tested during dopaminergic medication (‘on’) and dopaminergic withdrawal (‘off’). (n.s.: non-significant, s.: significant).

|  |  |  |  |
| --- | --- | --- | --- |
| 1 a |  |  |  |
|  | PD (N=15) | HCs (N=14) |  |
| | Mean ± SD | Mean ± SD | p (< 0.05) |
| Sex | 10 M, 5 F | 9 M, 5 F | n.s. (0.89) |
| Age | $69.93 \pm 8.75$ | $69.57 \pm 6.06$ | n.s. (0.90) |
| Education | $11.60 \pm 4$ | $13.07 \pm 2.58$ | n.s (0.25) |
| MMSE | $29.13 \pm 0.64$ | $29.07 \pm 1$ | n.s. (0.84) |
| MMPSE | $29.79 \pm 2.29$ | - | |
| 1b |  |  |  |
|  | PD on (N=15) | PD off (N=15) |  |
| | Mean ± SD | Mean ± SD | p (< 0.05) |
| UPDRS – III | $17.67 \pm 6.80$ | $37.21 \pm 10.04$ | s. (0.0001) |
| H&Y | $2.08 \pm 0.18$ | $2.32 \pm 0.32$ | s. (0.02) |

All participants were naïve as to the purposes of the study and signed the informed consent. The experimental protocol was approved by the local Ethics Committee at the IRCCS Santa Lucia Foundation of Rome (Reference number: CE/PROG.533) and was conducted in accordance with the ethical standards of the 2013 Declaration of Helsinki.

### Apparatus and Stimuli

Participants sat in a Cave Automatic Virtual Environment (CAVE) with projectors directed to four walls of a room-sized cube (3 m X 3 m X 2.5 m; Cruz-Neira, et al., 1993). The virtual scenario consisted of a basic room with a table (scale 1:1). At the center of the table, a dark yellow parallelepipedon was located with a blue glass on top of it. The virtual glass was placed in the participant’s peripersonal space at a distance of ∼ 50 cm. Participants observed from a first-person perspective (1PP) a virtual right arm, projected outside their right shoulder, and congruent in dimension and shape with their real body (see Fig 1. A). The virtual arm and the scenarios were created by means of Autodesk Maya 2015 and 3DS Max 2015, respectively. The kinematics of the avatar were realized in 3DS Max and implemented in the virtual scenario as an animated 3D mesh. The Virtual reality-EEG experiment was performed in an immersive three-dimensional immersive virtual environment rendered in CAVE by means of XVR 2.1 (Tecchia et al., 2014). Participants observed the virtual environment displayed through the Optoma 3D active glasses while their head position was tracked in real-time by means of an Optitrack System composed by eight infrared cameras placed inside the CAVE.

**Figure 1.**
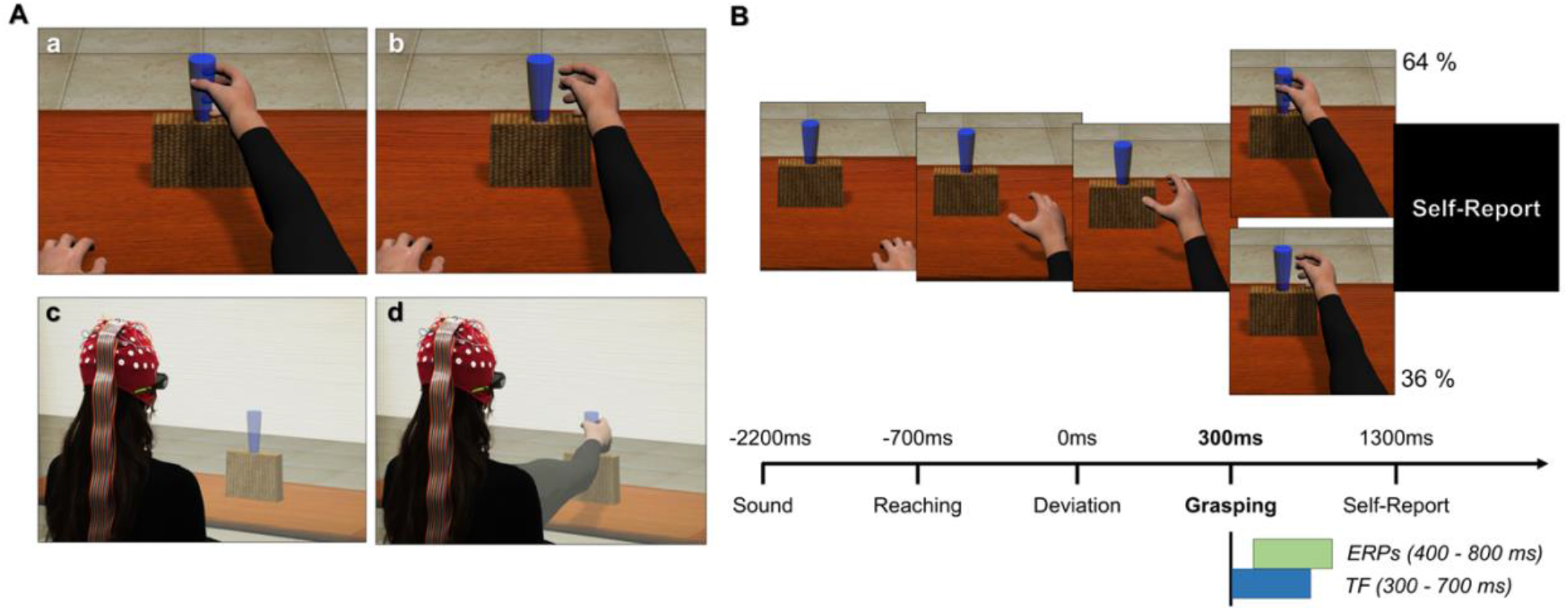
A: example of the experimental paradigm and setup. The image (c,d) shows the participant immersed in virtual scenario by the CAVE (cave automatic virtual environment) system, while observing the real-size virtual arm in first-person perspective, during a correct (a) or erroneous (b) grasping action. B: timeline of a single trial. The avatar’s action lasted ∼1,000 ms: the reaching phase was equal for both types of movements. The onset of the avatar’s arm-path deviation is set at 0 ms; the end of the avatar’s action occurs at 300 ms. The time windows for ERPs and TF analyses have been chosen a priori, based on existing literature.

### Experimental Procedure

Patients were tested in different sessions in separate days. First, an extensive neuropsychological assessment was administered while patients were under their dopaminergic treatment, to ascertain their cognitive profile, as part of clinical practice at the Foundation. For the experimental Virtual reality-EEG task, the patients visited the laboratory in two separate sessions, 15 days apart. In one session they were examined within 60 minutes from the first medication intake (‘on’ condition), while in the other after 18 hours washout from the individual prescriptions of dopaminergic medication used to treat PD (‘off’ condition; Langston et al., 1992). The order of on-off condition was counterbalanced across participants.

Before the beginning of the Virtual reality-EEG experiment, participants underwent a calibration phase where the size and the position of the virtual right arm was adapted to their real one. A short period of resting-state was included before beginning the experiment, which was not of interest for the current study. Then, they performed a brief practice session (8 trials, 4 correct and 4 erroneous) in which they familiarized with the virtual arm’s movements and the task. Each participant was requested to passively observe the virtual arm’s movements (by avoiding any real movements with their real upper limbs) and was informed that the goal of the movements was to reach and grasp the glass on the table. They were also informed that the action might or might not be successful. The Virtual reality-EEG task consisted in 110 trials per participant (70 correct and 40 incorrect virtual arm’s movements) with a total duration of ∼20 min. The choice of including a smaller proportion of error rather than correct actions was in line with the literature in which an error is often a salient infrequent event (Ullsperger et al., 2014). Previous studies with the current paradigm have used a proportion of 30-70% incorrect-correct actions (Pavone et al., 2016; Spinelli et al., 2018), as well as the inverted proportion of 70%-30% incorrect-correct events (Pezzetta et al., 2018). In this work, the choice to include a smaller number of errors (36%) compared to correct actions (64%) was preferred to keep the infrequency of occurrence of an error and at the same time guarantee a good number of trials, given the fact that a clinical and aging population was tested. At the onset of each trial, a sound signaled the beginning of the action. During the trial, participants passively observed from a first-person perspective the movement of the virtual right arm. The total duration of the movement was 1000 ms; the kinematics of the movement, identical for the 70% of the action duration in both correct and incorrect conditions, could diverge in movement’s trajectory in the last 30% of the time, leading to either a successful or unsuccessful grasp (Pavone et al., 2016; Pezzetta et al., 2018; Spinelli et al., 2018; Spinelli et al., 2022). The deviation from the to-be grasped object was identical in all the erroneous trials (Figure 1, panel B). The sequence of correct and incorrect trials was pseudorandomized. After the end of the action, the avatar’s arm remained still for 1000 ± 50 ms before a black screen appeared. During the inter-trial interval (ITI), one of three events occurred: 1) in 10 out of 110 trials (4 incorrect, 6 correct), participants had to answer a catch question “Did the arm grasp the glass?” (yes/no) in order to verify the engagement in the Virtual reality-EEG task; 2) in 65 out of 110 trials, an empty black screen was presented; and 3) in 35 out of 110 trials (13 incorrect, 22 correct), participants had to rate their illusory sense of embodiment over the virtual arm. The illusion was verbally rated on a visual analog scale (VAS) between 0 and 100 answering the question “To what extent did you feel the virtual arm was yours?” (0 = no ownership to 100 = maximal ownership; (Casula et al., 2021; Fusaro et al., 2019; Fusco et al., 2020; Pyasik, 2020; Tieri et al., 2015a,b, 2017). In the first and third type of event, the black screen lasted until a vocal response was given, whereas in the second event, the experimenter pressed a key to start the next trial, producing a variable ITI (mean duration: ∼ 4.000 ms, paradigm).

After each EEG session, an expert neurologist administered PD patients the UPDRS-Part III (Fahn & Elton, 1987; a 27 items scale where each item is evaluated on a 5-point Likert scale, ranging from 0 to 4) and the Hoehn and Yahr scale (H&Y, Hoehn & Yahr, 1967; this scale identifies 8 illness stages, indicated with the following numbers: 0-1-1.5-2-2.5-3-4-5). These scales (UPDRS III and H&Y) estimate the patients’ motor performance and allows to evaluate the efficacy of the dopaminergic medication in improving motor symptoms (higher scores mean higher disease severity). The two scales were administered in both ‘on’ and ‘off’ medication condition.

#### EEG recording and processing

EEG signals were recorded using a Neuroscan SynAmps RT amplifier system and 62 scalp electrodes embedded in a fabric cap (Electro-Cap International), arranged according to the international 10–10 system. Horizontal electro-oculogram was recorded bipolarly from electrodes placed on the outer canthi of each eye. Online, EEG signal was recorded continuously in alternating current mode with a bandpass filter (0.05–200 Hz) and sampling rates of 1.000 Hz. Impedances were kept under 5 kΩ. All electrodes were physically referenced to an electrode placed on the right earlobe and re-referenced offline to the common average across all electrodes. Offline, raw data were band-pass filtered with a 0.1-100 Hz filter (finite impulse response filter, transition 40–42 Hz, stopband attenuation 60 dB). Independent component analysis (ICA; Jung et al., 2000) was performed on the continuous EEG signal and components that were clearly related to blinks and ocular artifacts were removed (on average, 5.8 ICA components). For ERP analyses, an additional bandpass filter (0.3–30 Hz) was applied on the continuous raw signal. EEG signal was then down-sampled to 500 Hz and epoched in wide windows of 3-s length, from -1.5 to +1.5 s to avoid edge artifacts induced by the wavelet convolution in the time-frequency analysis. Epochs were time-locked (0ms) at the avatar’s arm-path deviation, with DC offset correction to the previous 300 ms preceding the deviation (Moreau et al., 2020; Pezzetta et al., 2018). Each epoch was then visually inspected to remove residual artefacts (e.g. eye blinks) by checking for epochs exceeding ±100 µV amplitude, (Drisdelle, Aubin, & Jolicoeur, 2017). After this procedure, a low number of trials was rejected from the original datasets (HCs: 4.5%, PD Dopa-ON: 1%, PD Dopa-OFF: 4%). Therefore, each group had a sufficient and comparable number of trials (mean ± SD; HCs_TOTAL_: 105 ± 5; HCs_COR_= 67 ± 3, HCs_INC_= 38 ± 2. PD Dopa-ON_TOTAL_: 109 ± 3, PD ON_COR_=69 ± 4; PD ON_INC_= 39 ± 1; PD Dopa-OFF_TOTAL_: 106 ± 5. PD Dopa-OFF_COR_=67 ± 3, PD Dopa-OFF_INC_= 39 ± 2; Pontifex et al., 2010).

Unless otherwise specified, data were normally distributed (Shapiro-Wilk test); thus, parametric analyses were adopted. Analyses were performed using the Brainstorm toolbox (free open source for MEG/EEG analysis, https://neuroimage.usc.edu/brainstorm/; Tadel et al., 2011) and customized Matlab routines. Statistical analyses were performed using R software (R Core Team 2014). Effect sizes were calculated using Cohen d formula. ERPs and time-frequency statistical analyses were made using the *erpR* package (Arcara & Petrova 2014). Practice trials were excluded from the analyses.

#### EEG analyses

##### Analysis in the Time Domain

All ERPs analyses were based on mean amplitude, as recommended by (Luck, 2005). To analyze ERPs at a whole brain level, we performed a time-point cluster-based permutation analyses with 1000 repetitions (with significant differences for clusters set for p < 0.05) and MonteCarlo correction in the 0-1000 ms time window on all electrodes, with cluster comparison within and between groups, which is appropriate to assess the reliability and robustness of neural patterns over neighboring data points (Formica et al., 2021; <u>Maris & Oostenveld, 2007</u>). Also, analyzing all electrodes allow to capture potential data-driven modulations on all electrode level which can be of relevance in aging or clinical populations, where shifts in neural activation and compensatory processes might be observed (van Dinteren et al., 2018). In addition, traditional ERPs analyses were performed. oERN was not analyzed as it was not found during visual inspection of the time series in the a priori selected time-window (see discussion). The oPe is a P300-like component maximally peaking at electrode Pz (Overbeek et al. 2005). Planned comparisons within groups on the variable “condition” were performed on a-priori established time-windows of interest (400-800 ms) on the electrode of interest (Pz) to be consistent with previous studies (Pavone et al., 2016; Pezzetta et al., 2018; Spinelli et al., 2018). Then, by following golden-standard recommendations (Luck, 2014; Kappenman & Luck, 2016), the ERP differential (obtained by subtracting the correct from the erroneous condition) was compared with three t-tests namely one within the PD “Group” (PD on vs PD off), and two across the HCs vs PD groups (HCs vs PD ‘on, and HCs vs PD ‘off’).

##### Analysis in the Time-Frequency domain

For the time-frequency analysis, we used a complex Morlet transformation to compute time-frequency decomposition. A mother wavelet with central frequency of 1 Hz and 3 s of time resolution (full width half maximum, FWHM) was designed as in Brainstorm software (Tadel et al., 2011). The other wavelets were computed from this mother wavelet and ranged from 1 to 80 Hz, with 0.5-Hz logarithmic frequency steps. To normalize each signal and frequency bin separately with respect to a baseline, we computed the relative power change (in %) over the time-frequency decomposition as

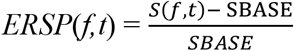

where S(t, f) is the signal spectrum at a certain given interval of time (t) and frequency (f), and S_base_ represents the mean power of the reference signal of the baseline interval (event related spectral perturbation, ERSP). To avoid edge effects, the power activity from -700 to - 500 ms - the window in which the avatar’s movement was identical in erroneous and correct conditions-was used as baseline interval (S_base_). Positive and negative values index a decrease or an increase in synchrony of the recorded neuronal population (Pfurtscheller, Neuper, Brunner, & Lopes Da Silva, 2005) with respect to a given reference interval, where equal neural activity is expected between conditions. In our case, a relative power increase/decrease represents a modulation of power compared with the mean power activity at baseline (Figure 4). To investigate the effect on the whole brain, we performed a time-point cluster-based permutation analyses with 1000 repetitions for each run (p < 0.05) and Montecarlo correction on a wide window from 0 ms to 1000 ms to see the distribution on the scalp. Cluster comparisons within and between groups were performed. Also, in line with previous studies (Moreau et al., 2020; Pavone et al., 2016; Pezzetta et al., 2018; Spinelli et al., 2018), the main analyses for theta activity were computed on the FCz electrode, focusing on theta band (4-8.1 Hz) in the preselected time interval (300-700 ms) corresponding to a total of 400 ms from the end of avatar’s action. For the analyses at the electrode level (FCz), for each group, planned comparisons with a single factor “Condition” with two levels (correct/erroneous) was performed. Then, the theta band differential (obtained by subtracting the erroneous from the correct condition) was compared with three separate analyses (Luck, 2014; Kappenman & Luck, 2016: one t-test in PD with the within factor “Group” with two levels (PD on/PD off), and two across group t-tests (HCs/PD on; HCs/PD off). This has been done to follow the methodology of the cluster-based permutation, based on t-test comparisons. Beside theta (4-8.1 Hz), analyses at a cluster level were performed also on the frequencies of potential interest for error monitoring processes in Parkinson’s disease namely: delta (2-4 Hz), alpha (8.1 –12.3 Hz), and beta (12.3–30.6 Hz) bands (Koelewijn et al., 2008; Luu et al., 2004; Moran et al., 2011).

### Clinical and neuropsychological testing

Clinical data ascertaining motor ability in relation to dopaminergic medication were analyzed. For UPDRS III and H&Y scales, two ANOVAs with dopaminergic “Medication” as factor with two levels (on/off) were performed. Correlations between clinical scales (UPDRS, H&Y) and EEG signals were performed to investigate clinical deficits in relation to EEG states during different dopaminergic conditions.

Neuropsychological tests assessing executive functions and cognitive control were also administered to the PD ‘on’, as previous data suggested a relation between theta/ern activity and performance in tasks underlying executive functions (Eckart et al., 2014; Sauseng et al., 2005), such as the Trial Making Test (B and BA) and the Wisconsin Card Sorting Test (Kim Myung et al., 2006; Willemssen et al., 2008). Correlations between theta activity and tests assessing executive functions were conducted. Details on Methods and Results concerning the neuropsychological data can be found in Supplementary Materials.

### Subjective reports

Embodiment ratings and the catch answers were calculated for correct and erroneous actions in the three groups as in previous studies (Pezzetta et al., 2018; Spinelli et al., 2018). To explore the link between sense of embodiment and electro-cortical indices of error processing, Spearman correlations between subjective reports and EEG signals were computed. Further details on Methods and Results for embodiment ratings can be found in Supplementary Materials.

Statistical analyses of clinical-neuropsychological data and subjective reports were performed using R software (R Core Team 2014). Greenhouse-Geisser correction for non-sphericity and Bonferroni correction for multiple comparisons were applied, when appropriate.

### Data availability

The data are available in the Open Science Framework (OSF) repository https://osf.io/z9rbu/.

## RESULTS

### Clinical deficits in relation to the dopamine states as inferred from UPDRS III and H&Y scales

Confirming the beneficial effect of dopamine assumption for extrapyramidal symptoms, the UPDRS III scores of patients with PD decreased significantly from the ‘off’ (M = 37.21, SD = 10.04) to the ‘on’ (M = 17.67, SD = 6.80) treatment condition (F_(1,13)_ = 29.14, p= 0.0001, n^2^_p_= 0.69). Changes of H&Y scale values in the different dopamine levels point at a similar effect with significant higher values in ‘off’ (M = 2.32, SD = 0.32) than in ‘on’ (M = 2.08, SD = 0.18) condition (F_(1,13)_ = 7.71, p= 0.02, n^2^_p_= 0.37. See Table 1b). One patient was excluded from the analysis because off condition evaluation was missing. No significant correlation was found between UPDRS, H&Y and EEG signals (theta, oPe).

### EEG

#### Time-Domain Analysis

##### Cluster-based statistics

We found significant differences in the within group analysis in all three groups, but with dissimilar spatial distribution. In the HCs a significant difference (p = 0.008, range 360-876 ms) was found between correct and erroneous actions; similarly, the cluster-based permutation revealed a difference between the two conditions both in PD ‘on’ (p = 0.002, range 380-1000 ms) and in PD ‘off’ (p = 0.008, range 300-634 ms). PD ‘off’ showed greater voltage in the fronto-central rather than parietal electrodes after the observation of erroneous actions, showing an increased activity that was limited in time (Figure 2). Cluster-comparisons between groups did not show significant differences in the window of 0-1000 ms.

**Figure 2.**
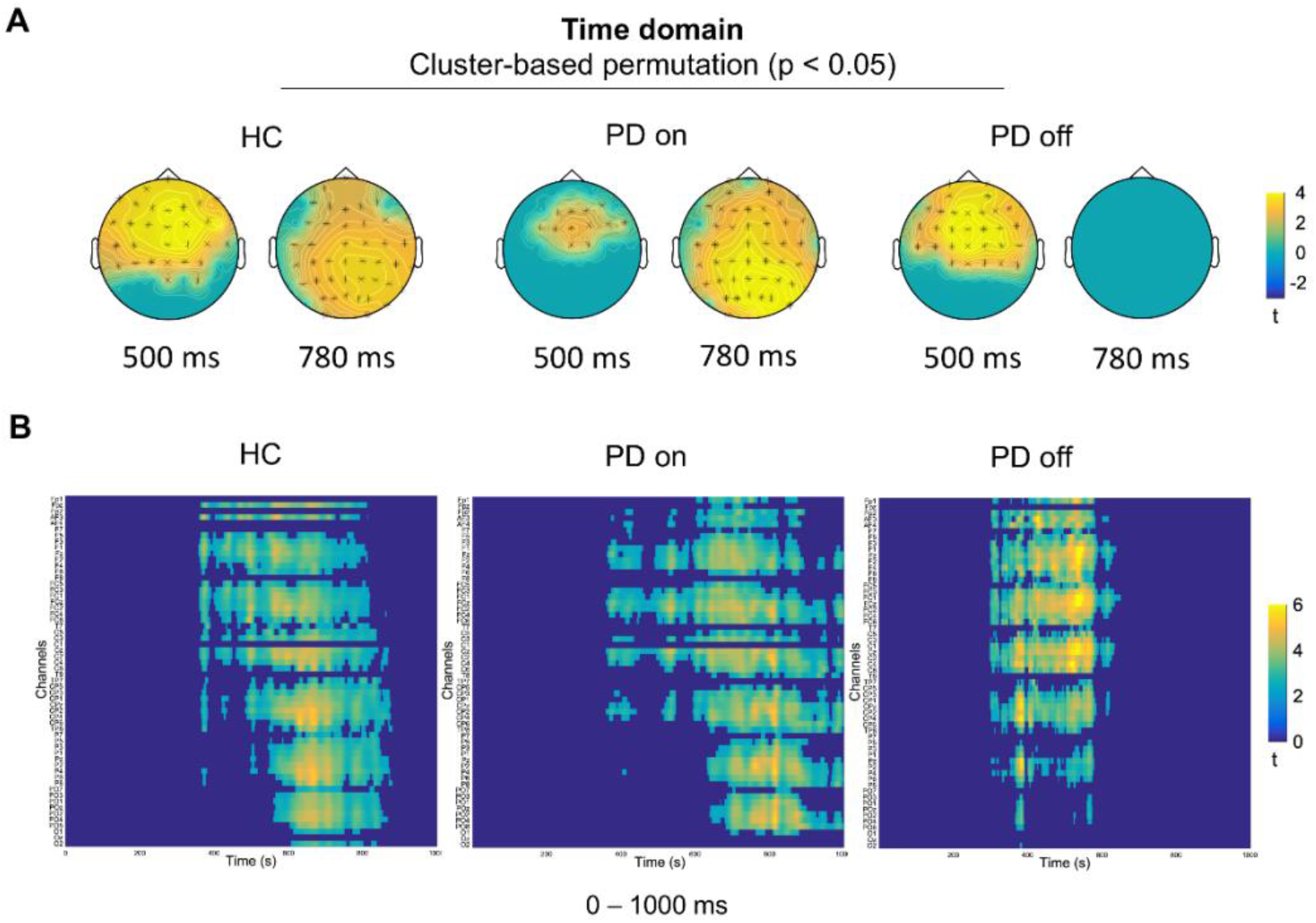
Cluster-based permutation in the time domain for each group. **A**. Scalp representation of the cluster-based permutations (dependent sample t-test with cluster-correction p<0.05) of erroneous versus correct action, extracted at two representative time points inside the window of interest. **B**. Channel (y-axis) x time (x-axis) representation of the cluster-based permutation for erroneous versus correct actions in the three groups.

##### Analyses on the electrode FCz

Analysis for the oERN were not performed as a clear peak was not found on visual inspection. For a possible interpretation of this negative finding, see the discussion section. **Analyses on the electrode Pz**. Traditional analyses on electrode Pz for the oPe (400-800 ms) showed that all groups had a significant difference between correct and erroneous actions (Figure 3); indeed HCs showed a significant difference, with greater amplitude for erroneous rather than correct actions [HCs: t(13)= -3.27, p = 0.006, d=0.65, M_ERR_= 7.59 µV, M_CORR_= 5.10 µV]; a significant difference was also found in the PD groups, both in ‘on’ [t(14)= -3.08, p = 0.008, d= 0.61; M_ERR_= 3.90 µV, M_CORR_= 2.06 µV] and ‘off’ [t(14)= -2.22, p = 0.04, d=0.40, M_ERR_= 4.96 µV; M_CORR_= 3.09 µV] condition, with greater oPe for erroneous than correct actions. Analyses of differential voltage (obtained by erroneous minus correct trials) between groups showed no difference, all groups had greater activity during erroneous trials, in the time window of interest. This result is in line also with the cluster-based permutation findings on all electrodes and on a wider window of time (0-1000), suggesting that all three groups (HCs, PD ‘on’ and PD ‘off’) showed a consistent result of greater amplitude in response to erroneous rather than correct actions.

**Figure 3.**
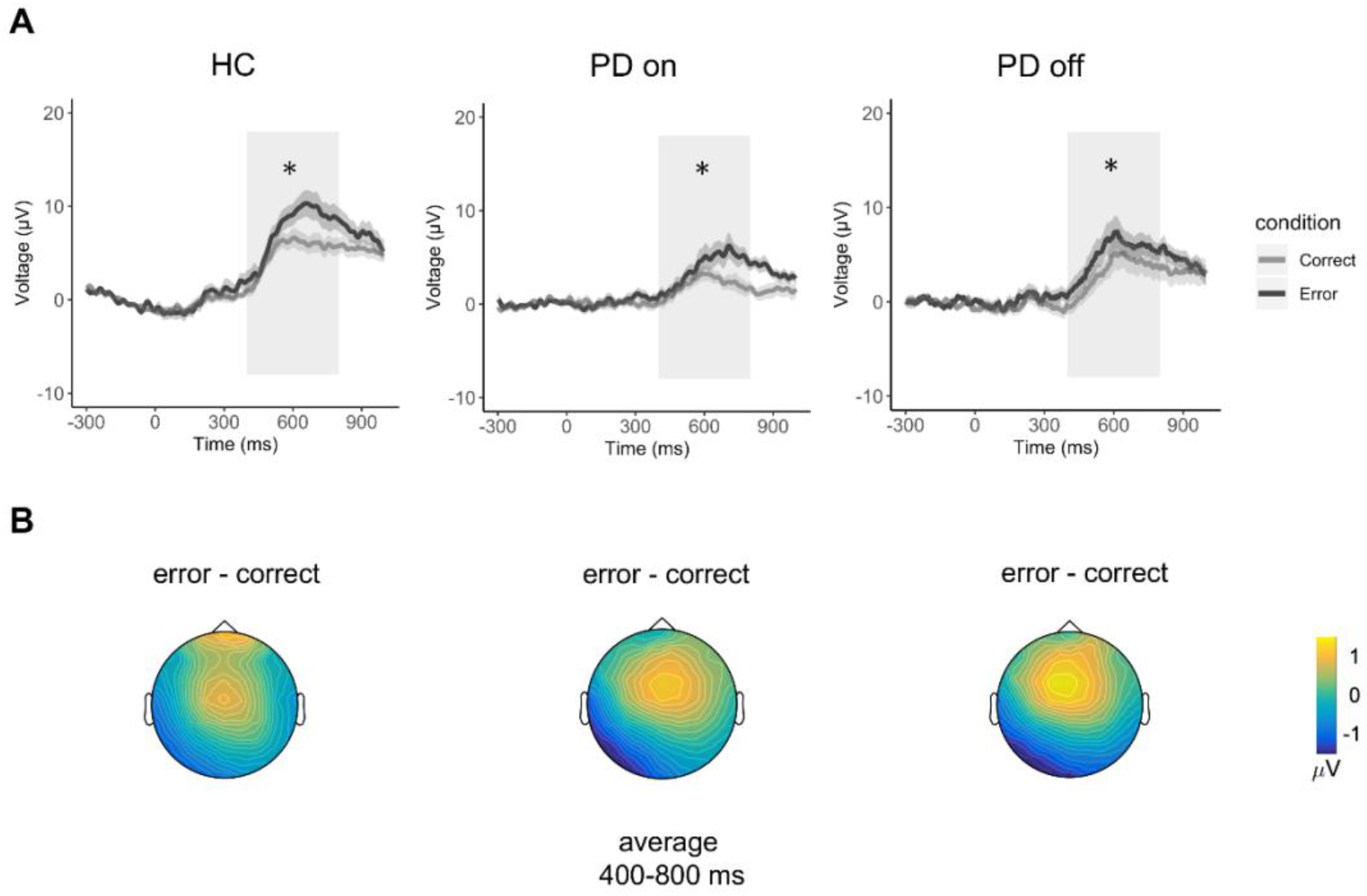
Electrophysiological results in the time domain for each group (ERPs). **A**. Grand average waveforms of oPe at electrode Pz. The end of avatar’s movement is set at 0ms. Lighter colors denote the standard error around the mean. The light-gray rectangle represents the interval window of analyses. **B**. Graphical representation of voltage distribution. The values are the result of the erroneous-minus correct actions.

### Time-Frequency Domain Analysis

#### Theta (4-8.1Hz)

##### Cluster-based statistics

We found a difference between erroneous and correct condition in the HCs (p= 0.004, range 208-888 ms) and in PD ‘on’ (p= 0.01, range 0-648 ms), with greater theta activity for erroneous actions; the difference was most pronounced over the central areas (see scalp distribution of the clusters, figure 6). In PD ‘off’ there was no significant error vs correct grasping difference. When the HCs and PD ‘off’ were compared, we found a significant difference (p = .01, range 392-792 ms), most pronounced in the frontal and posterior areas. No other significant difference between groups was found (Figure 4).

**Figure 4.**
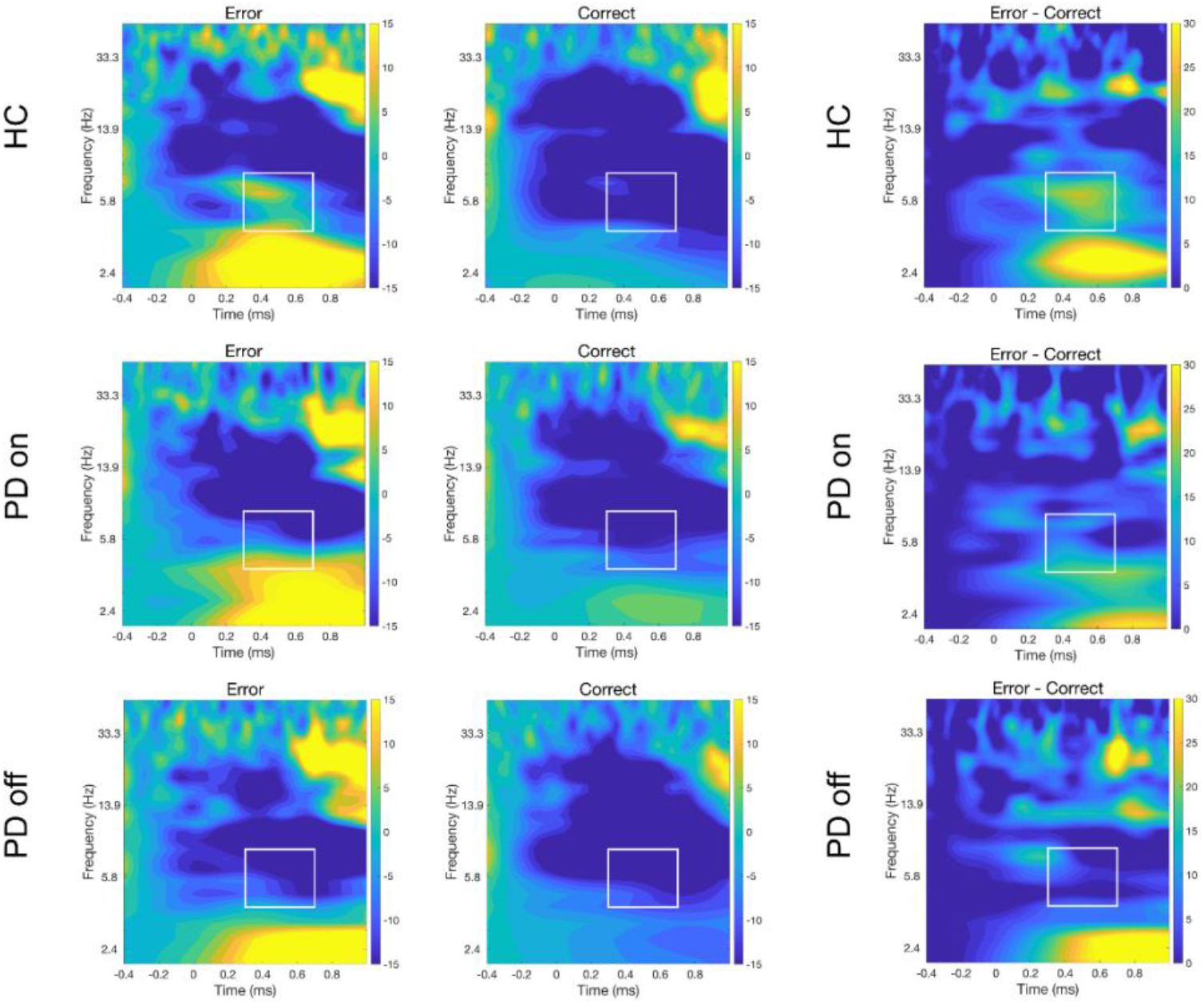
Time-frequency representation of Relative Power change (in %) with respect to the baseline for erroneous and correct conditions. The end of avatar’s arm-path deviation is set at 0ms. Erroneous and correct plots at electrode FCz in the three groups, frequencies from 1 to 50 Hz are displayed. In the third column, the differential plots are provided (erroneous – correct actions). The white rectangles highlight the a priori chosen window of interest between 300-700 ms and 4-8.1 Hz, that indicate the values used for statistical analyses.

##### Analyses on the electrode FCz

The HCs showed an effect of Condition [t(13)=-2.74, p=0.02, d=1.14, M_ERR_= 0.10, M_CORR_=-15.65], that was also present in PD ‘on’ [t(14)=-2.53, p=0.02, d=0.42, M_ERR_= -6.94, M_CORR_= -17.22], with greater theta activity for erroneous compared to correct actions. Contrary to the other two groups, ‘off’ patients did not show any difference [t(14)=0.68, p=0.51, d=0.14, M_ERR_= -17.08, M_CORR_= -19.74] (Figures 5, 6). Group comparisons on differential theta (erroneous – correct actions) showed a trend when the HCs and PD ‘off’ were compared [t(27)=1.90, p=0.067, d=0.71, M_HC_=15.75, M_OFF_=2.66]. We found no difference between the HCs and PD ‘on’: [t(27)=0.79, p=0.44, d=0.29, M_HC_=15.75, M_ON_=10.27] and PD ‘on’ vs. PD ‘off’: [t(14)=1.31, p=0.21, d=0.50, M_ON_=10.27, M_OFF_=2.66].

**Figure 5.**
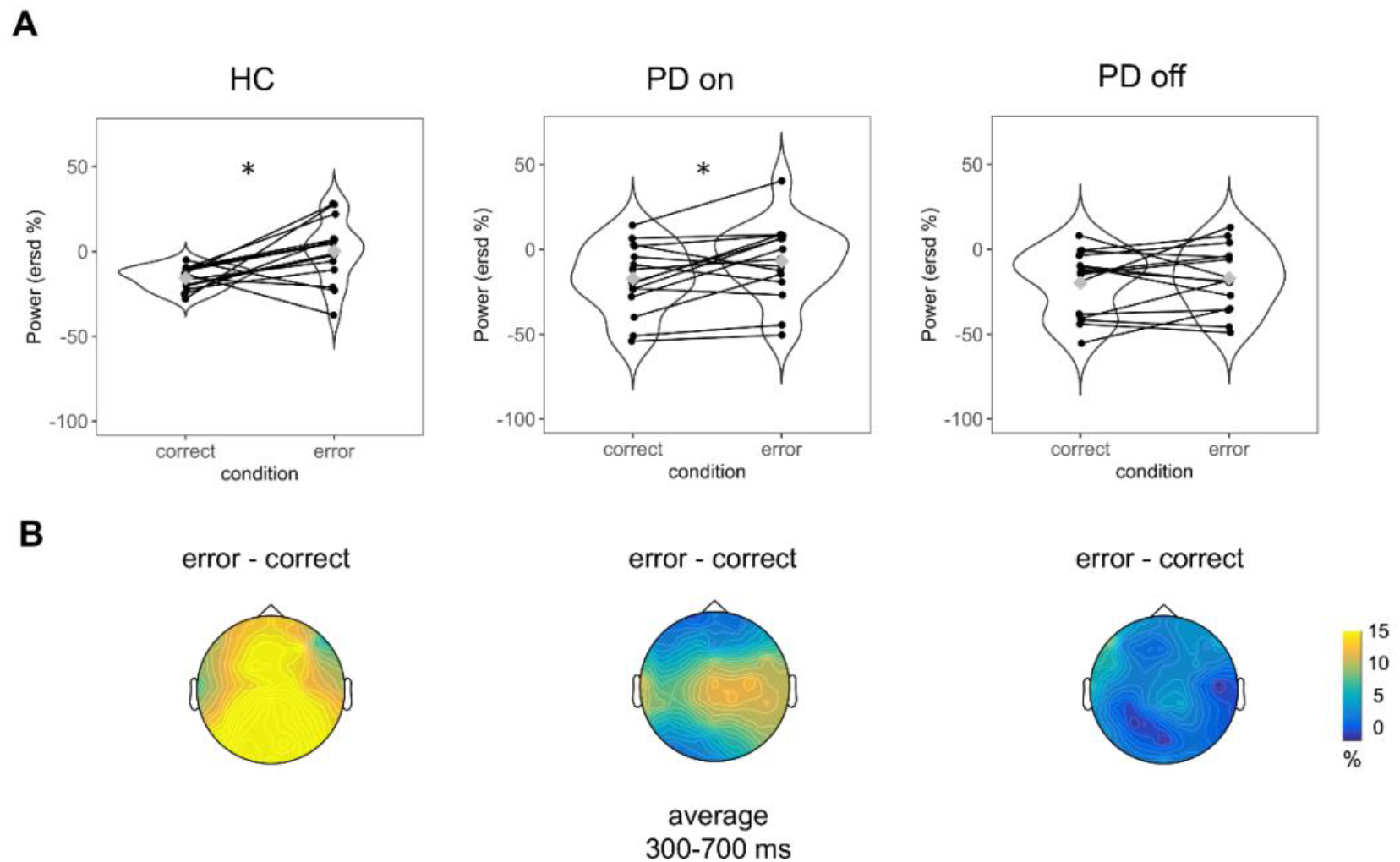
Graphical representation of Theta power (4-8.1 Hz) in the three groups. **A**. Violin plots represent theta activity in Correct and Erroneous actions. Y-axes represent theta power expressed in Relative Power change (in %). Gray diamonds in the violin plots represent the mean value; black lines connect individual subject observations (i.e., black points) in the two conditions. **B**. Graphical representation of voltage distribution. The values indicate the erroneous-minus correct action difference.

**Figure 6.**
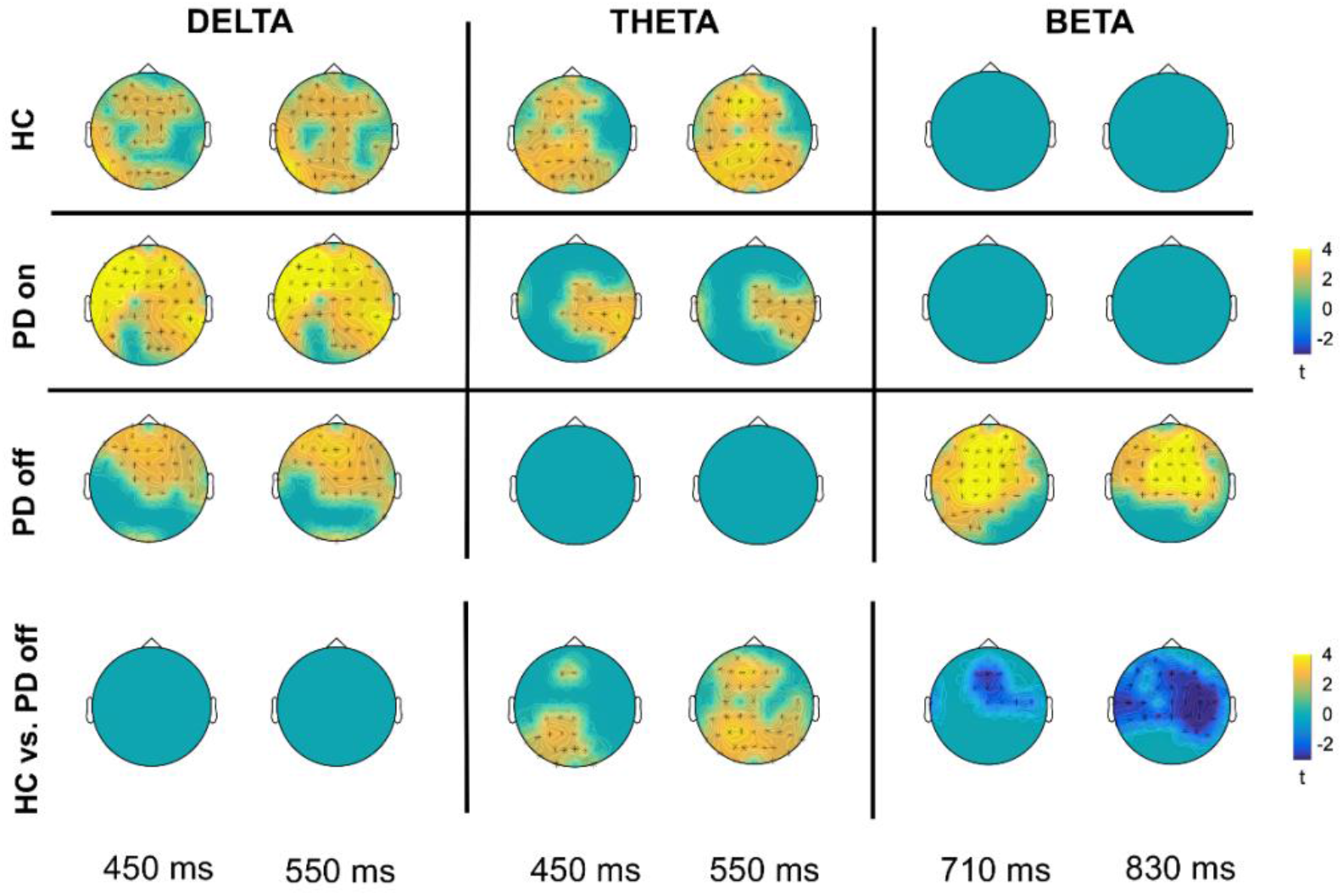
Cluster-based permutation in the time-frequency domain for each group. Scalp representation of the cluster-based permutation (dependent sample t-test with cluster-correction p<0.05) of erroneous versus correct action, extracted in two representative time points inside the window of interest. In the bottom line, cluster-based comparison between HCs and PD ‘off’ of differential activity (erroneous minus correct) in the frequency bands of interest. Analyses have been conducted on the wide window 0-1000 ms, two representative time points are shown.

### Other EEG frequencies potentially involved in error monitoring

#### Delta (2-4Hz)

##### Cluster-based statistics

We found significant difference for the three groups, respectively (HCs: p=0.008, range 0-1000 ms; PD ‘on’: p=0.002, range 0-1000 ms; PD ‘off’: p=0.004, range 0-1000 ms). The clusters showed greater delta activity for erroneous compared to correct actions (see Figure 6), in all three groups. In the HCs the difference was more prominent in the frontal and parietal areas, whereas in PD it was more prominent in the fronto-central areas. No statistical differences between groups (comparing the maps of erroneous – correct conditions) were found.

#### Alpha (8.1-12.3 Hz)

##### Cluster-based statistics

No significant activity was associated to erroneous rather than correct actions in any of the three groups.

#### Beta (12.3–30.6 Hz)

##### Cluster-based statistics

Cluster-based permutation indicates a trend in the HCs when erroneous and correct actions were compared (p= .07, range 280-440 ms) with central-contralateral distribution opposite to the observed arm (Figure 6). PD ‘on’ showed no significant difference of Condition. PD ‘off’ showed a significant difference (p = .004, range 150-1000 ms), mainly located on the central electrodes. The independent-samples comparison between groups revealed a significant difference between the HCs and PD ‘off’ (p=0.04, range 678-968 ms), accounted for by the fact that PD ‘off’ exhibited increased beta power in the central areas compared to the HCs (Figure 6, see Discussion).

## Discussion

To explore the influence of dopamine on the electrocortical dynamics of error monitoring we recorded EEG in patients affected by PD while they observed correct and erroneous actions performed by a virtual arm seen from a first-person perspective. Using a within-subject approach, the same patients were tested under dopaminergic medication (PD ‘on’) and after dopaminergic withdrawal (PD ‘off’). A control group of healthy participants was also included (HCs).

The first point of novelty is that an increase of theta power contingent upon observation of erroneous actions was found in HCs and PD ‘on’ but not in PD ‘off’ indicating that dopamine depletion modifies this neurophysiological marker of performance monitoring. The second one is that unlike theta activity, higher oPe and delta power for erroneous vs. correct actions was found both in HCs and PD patients (both ‘on’ and ‘off’ condition) indicating that error monitoring comprises distinct and independent neurophysiological processes that may or may not be impacted by dopamine balance.

### Dopamine does not modulate the oPe

Results revealed that observation of erroneous actions produced a clearly detectable oPe in all groups (electrode Pz). Importantly, however, cluster-based analyses showed that while HCs and PD ‘on’ displayed a distribution of activity spreading from frontal to posterior electrodes, PD ‘off’ had an effect that was mostly pronounced over fronto-central rather than parietal electrodes. This may be in keeping with studies showing that oPe is a cortical response characterized by subcomponents spreading over frontocentral and centroparietal electrodes (Overbeek et al., 2005) and could be associated with age-related and neurophysiological compensatory mechanisms (Iijima et al., 2000; Reuter et al., 2013). Notably, finding a similar oPe in PD ‘off’, PD ‘on’ and HCs, is consistent with the notion that generation of this component does not seem to depend on the dopaminergic system (Falkenstein et al., 2001). This evidence is in line with findings in other clinical populations, such as patients with ACC lesion (Maier et al., 2015; for systematic reviews see Lenzoni et al., 2021 and Pezzetta et al., 2021) in which the Pe – but not the ERN/theta - was unaltered. It was previously found that patients with lateral prefrontal lesion which extended to the insula had reduced Pe response (Ullsperger et al., 2002; Ullsperger & von Cramon, 2006), suggesting that the activity associated to the Pe activity might engage a neural network involving not only the ACC, but also anterior insula and somatosensory areas (Ullsperger et al., 2010), that may be spared in PD.

One may find surprising that we did not find any error-related negativity (oERN), a marker of error detection that was present in previous studies with young adults (Pavone et al., 2016; Pezzetta et al., 2018; Spinelli et al., 2018), but absent with older populations (Spinelli et al., 2022). We speculate that such absence may be related to weak modulation of oERN in aging (Nieuwenhuis et al., 2001; Thurm et al., 2020). Another non-alternative explanation is that time-frequency analyses might be better able to capture phase- and non-phase-locked activity during continuous actions (e.g. theta) whilst ERPs are more tuned to discrete events (e.g. ERN; Wang et al., 2020). Indeed, prior data suggest that ERN is dominated by phase-locking of intermittent theta-band (Trujillo & Allen, 2007), but that the observation of an error also elicits non-phase locked activity; thus, not all mid-frontal activity is associated to oERN generation (Moreau et al., 2020; Pezzetta et al., 2018).

### Dopamine does influence error-related, mid-frontal theta activity

PD in ‘off’ phase exhibited abnormal theta band activity with no power increase in response to errors. Crucially, when the same PD were tested just after their regular assumption of dopaminergic medication (PD ‘on’), theta activity in response to errors was restored leading to the same HCs pattern (Cavanagh & Frank, 2014). One may note that, similarly to previous studies (Singh et al., 2018; Willemssen et al., 2008), we did not find a direct theta activity difference contingent upon error monitoring between PD ‘on’ and ‘off’. Importantly, however, a significant theta power difference in response to action errors between HCs and PD ‘off’ was found. Compared to previous studies on PD (Seer et al., 2017), our patients in ‘off’ condition had a long withdrawal phase (∼ 18 h) and increased severity of extrapyramidal motor symptoms (mean UPDRS = 37) from their dopaminergic medication, which might have allowed to better highlight the contribution of dopamine in performance monitoring, when compared with their ordinary pharmacological treatment, compared to previous reports in which no difference was reported (Holroyd et al., 2002; Singh et al., 2018). Tellingly, no differential error-related theta activity was found between HCs and PD ‘on’, further hinting at the central role of dopamine in performance monitoring-related mid-frontal theta and in regulating the precision of information during predictive processes, triggered by salient and unexpected events (Friston & Kiebel, 2009), as recently found also in social context (Moreau et al. 2022; Solié et al., 2022; Boukarras et al., 2022).

### Delta, alpha and beta frequencies in response to errors

Studies indicate that alpha (van Driel et al., 2012), delta (Luu et al., 2004; Ulsperger et al., 2014), and beta (Koelewijn et al., 2008) frequencies may be potentially associated to error monitoring processes. In the present research, no error related modulation was found for the alpha band. Instead, delta activity turned out to be higher for erroneous than correct actions in all groups. This is in keeping with the notion that in a filtered signal, delta activity is associated with the Pe response in the time-domain (Luu et al., 2004; Kolev et al., 2005; Ullsperger et al., 2014). Interestingly, results showed how this marker of error monitoring is not influenced by dopamine depletion. The fact that error-related delta activity – together with the oPe - was found enhanced in response to observed erroneous events in HCs and PD (both ‘on’ and ‘off’) suggest that this mechanism is preserved in PD, compared to the error-related theta response which was not found within the PD ‘off’, suggesting that dopaminergic projections may not have a direct prominent role with error-related delta/oPe generation.

Analysis of beta band showed error-related increased in PD off, within group and also when contrasted with HCs. Beta rhythm has been associated to sensorimotor control (Jurkiewicz et al., 2006; Pfurtscheller et al., 2005; Torrecillos et al., 2015), learning tasks (Viñales et al., 2021) and long-distance communication between visual and sensorimotor areas (Engel & Fries, 2010). Local field recordings from the subthalamic nucleus identified excessive beta activity in PD associated to with pathophysiological motor symptoms (Oswal et al., 2013), that was restored by the dopaminergic treatment (Doyle et al., 2005). Studies indicate that beta rebound was stronger after incorrect rather than correct actions, suggesting a potential role of beta in the evaluation of action significance and active response inhibition (Koelewijn et al., 2008). In our study, PD ‘off’ showed stronger error-related beta response. No such effect was found in the PD ‘on’ (in whom dopaminergic medication seem to suppress beta activity; Doyle et al., 2005). In sum, while HCs show greater involvement of theta rather than beta response to errors, PD ‘off’ seem to show an opposite pattern. Whether PD may compensate the involvement of mid-frontal theta with higher frequencies during dopaminergic withdrawal, has to be investigated in future studies. It may be of special interest to explore whether increased beta activity might be detrimental (Moran et al., 2011) or whether it might represent a compensatory mechanism rather than a pathophysiological marker (Pollok et al., 2013).

### Clinical deficits in relation to the dopamine states as inferred from UPDRS and H&Y scales

Concerning the relation between the UPDRS-H&Y and EEG signals, no significant correlation was found. On this regard, we can speculate that this might be due to the fact that the correlation was computed between a subjective measure (e.g. the UPDRS-H&Y scales are administer and rated by a clinician) and an objective measure (the EEG signal). Similar lack of correlation was found also in previous reports (Singh et al., 2018), in which patients in ‘on’ condition had lower scores than ‘off’ condition in the UPDRS-III, but this result did not correlate with the EEG signal. Further, the scores obtained with the UPDRS III are related to patients ‘motor ability and it allows to evaluate the efficacy of the dopaminergic medication in improving general motor symptoms; however, this might be not directly related with the error-monitoring signals obtained within this task, in which a direct and active motor performance was not required.

Our approach allowed to directly investigate how distinct electrocortical signatures to errors are differently affected by dopamine balance in PD. Among the strengths of the present study is that we used an ecological and short (∼ 20 minutes) task which, thanks to immersive virtual reality, made possible to test the brain reaction to errors in PD, without confounds due to movement speed or difficulty (Ozkan & Pezzetta, 2018; see Supplementary Materials for a deeper discussion on the embodiment results). We acknowledge, however, some potential limitations. One may observe that PD were tested twice, whereas the HCs only one. Yet, in keeping with previous studies (Singh et al., 2018) we can reasonably exclude a learning effect, because the adopted task involves simple action-observation, and it is not related to the acquisition of task-specific abilities. A second limitation is that other neurotransmitters might play a role in the performance monitoring, either through direct modulation of the ACC or by virtue of their influence on the DA system (Calabresi et al., 2006; Singh et al., 2018). Future studies should take into consideration the role of neurotransmitters like serotonin, norepinephrine, GABA and adenosine in cooperating with dopamine to orchestrate error processes (Jocham & Ullsperger, 2009) in samples with different phenotypes (Van Nuland et al., 2021).

In conclusion, we expanded research in old adults and neurological populations, by providing novel support to the idea that error-related signals (theta and oPe/delta) may reflect distinct structural, functional, and biochemical paths within the complex architecture of the performance monitoring system (Di Gregorio et al., 2018; Krigolson & Holroyd, 2007; Steinhauser & Yeung, 2010). The error-related modulation of theta activity contingent upon dopamine depletion reported in our study may pave the way to future studies on neurophysiological biomarkers related to prediction processing and model updating (Klein et al., 2007; Friston et al., 2012; Masina et al., 2022) that may ultimately help to understand higher-order cognitive control in Parkinson’s Disease.

## Paradigm video

https://www.youtube.com/watch?v=F-NEbOT1nh4

## Author Contributions

Conceptualization of the idea: R.P. and S.M.A. Patients recruitment and neuropsychological assessment: S.Z., S.T. Neurological assessment: A.P. Virtual reality implementation and setup, and video production: G.T. Data collection: R.P., D.G.O., V.E., G.T. Data analysis and figures: R.P. Data interpretation: R.P., V.E., S.M.A. Writing the original draft: R.P., S.M.A. Revision of the manuscript and final approval: all the Authors. Supervision of the project: A.C., C.C., S.M.A.

