## Supplementary Materials for "Selective influence of dopamine on electrocortical signatures of error monitoring: a combined EEG and immersive virtual reality study in Parkinson’s disease"

The present file contains the supplementary material of the article

### METHODS

#### Neuropsychological tests assessing executive functions

PD patients received an extensive neuropsychological assessment during the 'on' condition, as part of a specialized hospital clinical practice. In Supplementary Table S1 are shown demographic and clinicals information of PD patients and main neuropsychological tests that have been considered of interest for the current study.

| Subject | Age | Education | Illness duration (months) | L_Dopa equivalent | BDI | MMSE | MMPSE | TMT_A | TMT_B | TMTB-A | WCSTcat | WCST_p | WCST_np | MMSE_pc | MMPSE_pc | TMT_A_pc | TMT_B_pc | TMTB-A_pc | WCSTcat_pc | WCST_p_pc | WCST_np_pc |
|---|---|---|---|---|---|---|---|---|---|---|---|---|---|---|---|---|---|---|---|---|---|
| P01 | 83 | 8 | 264 | 870 | 0 | 29 | 31 | 61 | 131 | 70 | 6 | 1 | 0 | 28.7 | 32.57 | 29.05 | 26.29 | 0 | 6 | 0 | 0 |
| P02 | 58 | 13 | 108 | 650 | 11 | 29 | 31 | 30 | 72 | 42 | 6 | 2 | 1 | 26.2 | 30.07 | 13.95 | 16.44 | 2.47 | 6 | 1.83 | 0.5 |
| P03 | 77 | 13 | 84 | 750 | 5 | 29 | 28 | 51 | 99 | 48 | 6 | 1 | 0 | 27.3 | 28.38 | 30.23 | 37.48 | 7.21 | 6 | 0 | 0 |
| P04 | 58 | 8 | 276 | 810 | 12 | 28 | 31 | 110 | 265 | 155 | 6 | 0 | 3 | 26 | 30.94 | 94.1 | 209.91 | 115.79 | 6 | 0 | 2.51 |
| P05 | 72 | 13 | 156 | 600 | 4 | 29 | 29 | 36 | 133 | 97 | 6 | 0 | 0 | 26.7 | 29.05 | 19 | 83.11 | 64.09 | 6 | 0 | 0 |
| P06 | 70 | 13 | 84 | 550 | 5 | 30 | 31 | 51 | 117 | 66 | 6 | 0 | 1 | 30 | 31.05 | 34.49 | 68.65 | 34.13 | 6 | 0 | 0.48 |
| P07 | 78 | 5 | 204 | 400 | 13 | 29 | 25 | 86 | 326 | 240 | 4 | 5 | 9 | 28.7 | 26.77 | 52.78 | 212.17 | 159.36 | 4.32 | 3.05 | 7.56 |
| P08 | 82 | 8 | 24 | 312,5 | 0 | 29 | 25 | 55 | 127 | 72 | 4 | 3 | 13 | 28.7 | 26.57 | 23.77 | 24.5 | 0.69 | 4.38 | 0.49 | 11.79 |
| P09 | 72 | 13 | 30 | 425 | 3 | 30 | 30 | 46 | 81 | 35 | 3 | 3 | 12 | 30 | 30.05 | 9.62 | 0 | 0 | 3.21 | 1.83 | 10.76 |
| P10 | 60 | 13 | 84 | 725 | 7 | 30 | 31 | 20 | 77 | 57 | 4 | 5 | 15 | 30 | 30.4 | 10.27 | 49.6 | 39.31 | 4.06 | 4.75 | 14.79 |
| P11 | 68 | 5 | 24 | 750 | 21 | 29 | 32 | 61 | 246 | 185 | 5 | 3 | 8 | 27.9 | 32 | 34.14 | 151.84 | 117.68 | 5.17 | 2.16 | 6.86 |
| P12 | 57 | 18 | 36 | 400 | 6 | 28 | 31 | 19 | 74 | 55 | 4 | 6 | 10 | 25.2 | 29.38 | 18.38 | 83.4 | 65 | 4.04 | 5.89 | 10.22 |
| P13 | 79 | 13 | 288 | 650 | 7 | 30 | NA | 55 | 137 | 82 | 6 | 1 | 7 | 30 | NA | 33.27 | 72.51 | 39.2 | 6 | 0 | 6.23 |
| P14 | 68 | 18 | 144 | 950 | 18 | 29 | 32 | 30 | 83 | 53 | 3 | 4 | 8 | 26.2 | 32 | 22.1 | 69.89 | 47.76 | 3.17 | 3.15 | 7.88 |
| P15 | 64 | 13 | 156 | 650 | 5 | 29 | 30 | 43 | 104 | 61 | 6 | 0 | 1 | 26.2 | 29.4 | 31.53 | 102.14 | 70.58 | 6 | 0 | 0.67 |
| | | | | | | | | | | | | | | cutoff: | | | | | | | |
| | | | | | | | | | | | | | | 23.8 | 22.85 | 94 | 283 | 187 | 4.25 | 7.65 | 10.75 |

**Supplementary Table S1.** Demographic, clinical and a subset of neuropsychological tests which tap executive functions. PD Patients were tested during 'on' condition. The column labeled "L-Dopa equivalent" reports the daily dose of dopamine or a dopamine agonist taken by each patient. On the left side of the table demographic and clinical data are shown; in the central section of the table, raw scores for each patient are reported, while on the right-side of the table, the corrected values are shown ("pc" stands for post correction for age-education). Cutoff scores for each neuropsychological test are also reported at the end of the table. MMSE: mini-mental state examination; MMPSE: Mini-mental Parkinson State Examination; BDI: Beck depression inventory; TMT_A: trial-making test A; TMT_B: trial-making test B;

TMT_B-A: trial making test BA; WCST_CAT: Wisconsin Card Sorting Test_categories; WCST_P: Wisconsin Card Sorting Test_perseverative errors; WCST_NP: Wisconsin Card Sorting Test_non perseverative errors.

We reasoned that, changes of fronto-central theta may be related to executive and working memory functions, as previous studies suggested (Eckart et al., 2014; Fusco et al., 2018). Thus, we selected two clinical tests that supposedly tap the same functions and may be of interest in relation to error monitoring aspects in the PD population (Kim Myung et al., 2006; Willemssen et al., 2008), namely: Trial Making Test (TMT subtest A, Giovagnoli et al, 1996), Trial Making Test (TMT subtest B, Giovagnoli et al, 1996), Modified Card Sorting Test (MCST, Nocentini et al, 2002, with the MCSTcat for categories, MCST_p for perseverative errors and MCST_np for non perservative errors). It was also calculated the composite score of the TMT (i.e. the score obtained for the TMT B minus the score obtained at the TMT A: TMT-BA), which is traditionally computed to derive measures that highlight executive functions abilities (Sánchez-Cubillo et al., 2009). To assess the general cognitive functioning also the Mini Mental State Examination (MMSE; Measso et al., 1993) and Mini Mental Parkinson State Examination (MMPSE; Costa et al., 2013) were administered. We performed correlations between all-brain cluster-based permutation across time (0-1000 ms) in theta activity and the executive functions tests, with Montecarlo correction for multiple comparison, to observe the scalp distribution across electrodes.

**Subjective reports during the Virtual reality-EEG task**

The embodiment question ("How much did you feel that the arm was yours" on a scale 0-100) was present only in a subset of trials (12 % of incorrect, 20 % of correct trials). For each participant, mean embodiment ratings for each type of trial were calculated. The scores were entered into three separate ANOVA with factors "Condition" (correct/erroneous) and "Group" (two anova with ‘between variable’: HCs-PD ‘on’; HCs-PD ‘off’; one anova with ‘within variable’: PD’on’-PD’off’). In order to explore the link between sense of embodiment and electro-cortical indices of error processing, Spearman correlations between Embodiment ratings and error signatures (Theta and oPe) were conducted. For the catch trials, the percentage of accuracy for each group was calculated.

### RESULTS

#### Correlations between theta activity and executive function abilities

Results show significant correlations in the PD ‘on’ group only, which is also the condition during which the patients received the neuropsychological assessment. Correlations between raw scores of neuropsychological tests and EEG activity were found significant between theta activity and TMT-B (only a trend level; $R=0.51$, $p=0.05$) and between theta activity and TMT BA ($R=0.53$, $p=0.04$). Clusters analyses showed significant clusters between theta and TMT B ($p= 0.02$; range 20-1000 ms) and theta and TMT BA ($p= 0.02$; range 20-1000 ms). No other significant correlations with executive functions’ tests were obtained.

**S1**

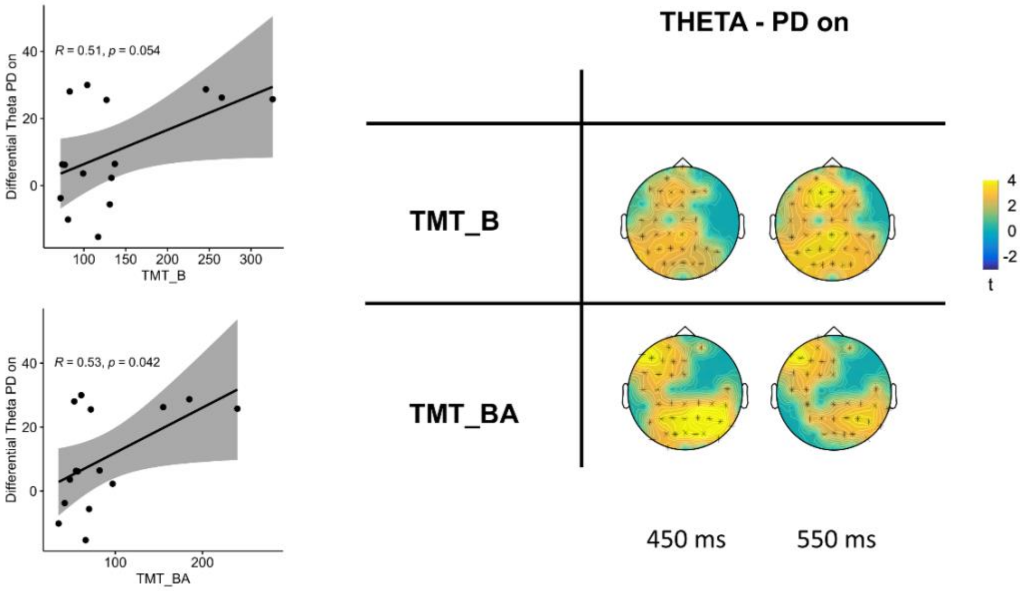

**Figure S1.** On the left side of the graph, correlations between theta activity in the time-window of 0-1000 ms and TMT_B (picture top-left) and TMT_BA (picture bottom-left). On the right, cluster-based permutation ($p<0.05$) between theta (4-8.1 Hz) in PD ‘on’ group and TMT-A and TMT-BA. Only effects that survived Montecarlo cluster-based multiple correction are showed.

##### Subjective reports during the Virtual reality-EEG task

Lower sense of ownership was reported for error than correct trials in all the three groups (HCs: $M_{ERR}$= 61.62, $M_{CORR}$= 62.55; PD ‘on’: $M_{ERR}$= 59.60, $M_{CORR}$= 65.37; PD ‘off’: $M_{ERR}$= 58.75, $M_{CORR}$= 63.77). However, analyses within and between groups showed no significant difference of ‘group’ or ‘condition’ (all ps > 0.05). Concerning the catch questions, overall participants in the three groups responded correctly to the questions, confirming the engagement in the task and the understanding of the observed action (correct answers: HC: 94%, PD ‘on’: 97%; PD ‘off’: 93%).

#### DISCUSSION

##### Correlations between theta activity and neuropsychological tests in PD

In the PD ‘on’ group only, we found a positive correlation between theta activity (differential score) and both TMT-B and the TMT-BA. In specific greater theta correlated with longer RTs to perform the TMT task. While the TMT A requires mostly visuoperceptual abilities, the TMT B reflects primarily working memory and task-switching abilities. Tellingly, the TMT-BA provides an indication of executive control abilities (Sánchez-Cubillo et al., 2009). The fact that theta activity in PD ‘on’ did not correlate with the TMT A (only visuoperceptual abilities) but did correlate with the TMT B and TMT BA might indicate a link between fronto-central slow-frequencies and increased cognitive effort. Previous studies have also found a link between fronto-parietal theta activity and executive functions, such as working memory (Sauseng et al., 2005) and eye-blink decay (Chen et al., 2016), which have been associated with executive dysfunction in PD. Even though this might suggest a possible link between midfrontal error-related theta and executive functions, it is to be highlighted that the reported correlations were not corrected for multiple comparison and deserve future investigations.

**Effect of dopamine on subjective performance in the Virtual reality-EEG task**

Concerning the subjective reports, even if qualitatively participants reported greater sense of ownership during the performance of a correct rather than erroneous action, analyses of the embodiment ratings did not show a significant difference between conditions. This result is at odds with respect to previous studies in young adults (Pavone et al., 2016; Spinelli et al., 2018; Pezzetta et al., 2018). Although somewhat speculative, several explanations for the across-studies differences can be offered. The first one is that, at variance with our previous reports, here we tested old adults. Another difference is that, in this study, we asked participants to report embodiment over the artificial limb in a 30% (rather than in 100%; Pavone et al., 2016) and we only asked questions concerning the feeling of ownership and not the feeling of agency, which has been also linked to action monitoring (Villa et al., 2018, 2021), to avoid long sessions for patients' fatigue.

Concerning the catch questions, PD mostly answered the questions correctly. We indeed expected a high rate of accuracy as the assignment was extremely easy, and it was not to be performed under time pressure conditions. Beside the groups not showing an effect of embodiment in the current experiment, previous studies suggested the advantage of using a virtual task to induce the illusion of committing an error in 1PP (Pavone et al., 2016), in which correct/incorrect actions are performed by an arm calibrated on the participants' body size, something that would be impossible to adapt with traditional setups; on this line, many studies are recently suggesting how the VR could be a promising technique for cognitive and motor rehabilitation (Tieri et al., 2018).
